# Differential Synaptic Input to External Globus Pallidus Neuronal Subpopulations *In Vivo*

**DOI:** 10.1101/2020.02.27.967869

**Authors:** Maya Ketzef, Gilad Silberberg

## Abstract

The rodent external Globus Pallidus (GPe) contains two main neuronal subpopulations, prototypic and arkypallidal cells, which differ in their cellular properties. Their functional synaptic connectivity is, however, largely unknown. Here, we studied the membrane properties and synaptic inputs to these subpopulations in the mouse GPe. We obtained in vivo whole-cell recordings from identified GPe neurons and used optogenetic stimulation to dissect their afferent inputs from the striatum and subthalamic nucleus (STN). All GPe neurons received barrages of excitatory and inhibitory input during slow wave activity. The modulation of their activity was cell-type specific and shaped by their respective membrane properties and afferent inputs. Both GPe subpopulations received synaptic input from STN and striatal projection neurons (MSNs). STN and indirect pathway MSNs strongly targeted prototypic cells while direct pathway MSNs selectively inhibited arkypallidal cells. We show that GPe subtypes are differently embedded in the basal ganglia network, supporting distinct functional roles.

## Introduction

The Globus pallidus *pars externa* (GPe, also called the GP in rodents) is a central part of the basal ganglia (BG). The GPe is composed of continuously active GABAergic neurons and is traditionally considered part of the indirect pathway of the BG (Albin et al., 1989; DeLong, 1990). Within this framework, the GPe mainly projects to the output structures of the BG, the substantia nigra *pars reticulata* (SNr) and the internal segment of the globus pallidus (GPi, also referred to as entopeduncular nucleus in rodents), as well as the subthalamic nucleus (STN). The cardinal input to the GPe is GABAergic inhibition from striatopallidal medium spiny neurons (MSNs) of the indirect pathway, however, it is also reciprocally connected to the STN (Kita et al., 1983; Kita and Kitai, 1991; Robledo and Feger, 1990) and receives input from axon collaterals of striatonigral MSNs (Cazorla et al., 2014; Kawaguchi et al., 1990; Wu et al., 2000). Thus, the GPe is positioned as a hub, which connects the main BG pathways: the direct-, indirect-, and hyper-direct pathways (Mathai and Smith, 2011; Nambu et al., 2002). Additionally, the GPe was shown to project to the cortex and thalamus (Mastro et al., 2014; Saunders et al., 2015). This suggests that rather than acting as a relay station, the GPe plays a far more central role in BG function than previously appreciated.

The GPe was traditionally regarded as a homogenous nucleus, yet already in early studies (DeLong, 1971), different firing patterns of GPe neurons were described. It is now clear that GPe neurons are divided into at least two subpopulations, with distinct electrophysiological properties, developmental origins, molecular markers and projection targets (Abdi et al., 2015; Dodson et al., 2015; Hernandez et al., 2015; Mallet et al., 2012; Mastro et al., 2014). The majority of GPe neurons, namely prototypic cells, conform to the classic mode; they are high frequency spiking neurons that project to downstream nuclei, with a small portion of them also projecting to the striatum (Bevan et al., 1998; Dodson et al., 2015; Mallet et al., 2012; Mastro et al., 2014; Saunders et al., 2016). Prototypic cells express the transcription factor NK2 homeobox 1 (NKX2.1) (Abdi et al., 2015; Dodson et al., 2015) and to a lesser degree also parvalbumin (PV) and LIM homeobox 6 (LHX6) (Abdi et al., 2015; Dodson et al., 2015; Mallet et al., 2012; Mastro et al., 2014; Mastro et al., 2017). The other subtype of GPe neurons consists of the arkypallidal cells, which are a smaller group of relatively less active neurons that project exclusively to the striatum (Mallet et al., 2012). These neurons express the transcription factor forkhead box protein P2 (FoxP2) (Abdi et al., 2015; Dodson et al., 2015), the neuropeptide precursor preproenkephalin (PPE) (Hoover and Marshall, 2002; Mallet et al., 2012) and the neuronal PAS domain protein 1 (NPas1) (Hernandez et al., 2015).

The marked differences between the two main subtypes of GPe neurons suggest that they have different functional roles, which would be further supported by distinct sets of afferent synaptic inputs on top of differences in membrane properties that were mostly described ex vivo thus far. The functional organization of synaptic inputs to the two different GPe subtypes is, however, largely unknown. In order to accurately position the GPe within the BG functional scheme, it is imperative to characterize the afferent synaptic inputs to the different subtypes of GPe neurons. Here, we studied the GPe using *in vivo* whole-cell recordings, enabling us to characterize the membrane properties, synaptic input, and activity of identified prototypic and arkypallidal cells in the intact brain. We quantified the afferent inputs to the respective cell types from STN and striatum, and also showed intra-pallidal connectivity between the two cell types.

## Results

Whole-cell recordings were obtained from neurons in the GPe of anesthetized mice (figure 1A-B), and indicated the existence of at least two main GPe neuronal subtypes. Most recorded neurons were spontaneously active with high firing rates (Figure 1 C and F), had depolarized membrane potential values (Figure 1 C and E), and paused or reduced their firing rate during cortical slow wave “up-states” (Figure 1 C). These neurons were further classified as prototypic GPe cells using two independent measures. Classification was done either post hoc by immunostaining for FoxP2 (Figure 1 C, left), or during recordings using the “*optopatcher”* (Katz et al., 2013; Ketzef et al., 2017) in Channelrhodopsin (ChR2) expressing cells of NKX2.1-ChR2 (17 cells from 15 mice) or PV-ChR2 mice (3 cells in 3 mice) (Figure S1). A smaller fraction of recorded neurons were silent (see example in S3) or fired at lower average frequencies (Figure 1 D and F), had more hyperpolarized membrane potential (Figure 1 D and E), and depolarized during cortical up-states (Figure 1 D). These neurons were identified post hoc as arkypallidal cells as they expressed FoxP2 (Figure 1 D, left). These activity profiles are in agreement with previous descriptions using extracellular recording methods (Abdi et al., 2015; Dodson et al., 2015; Mallet et al., 2012) and suggest that during cortical up-states, arkypallidal cells are largely excited while prototypic cells are inhibited. These results point towards a clear distinction between the two main GPe subpopulations recorded *in vivo,* which could reflect differences in their synaptic inputs and membrane properties.

**Figure 1:**
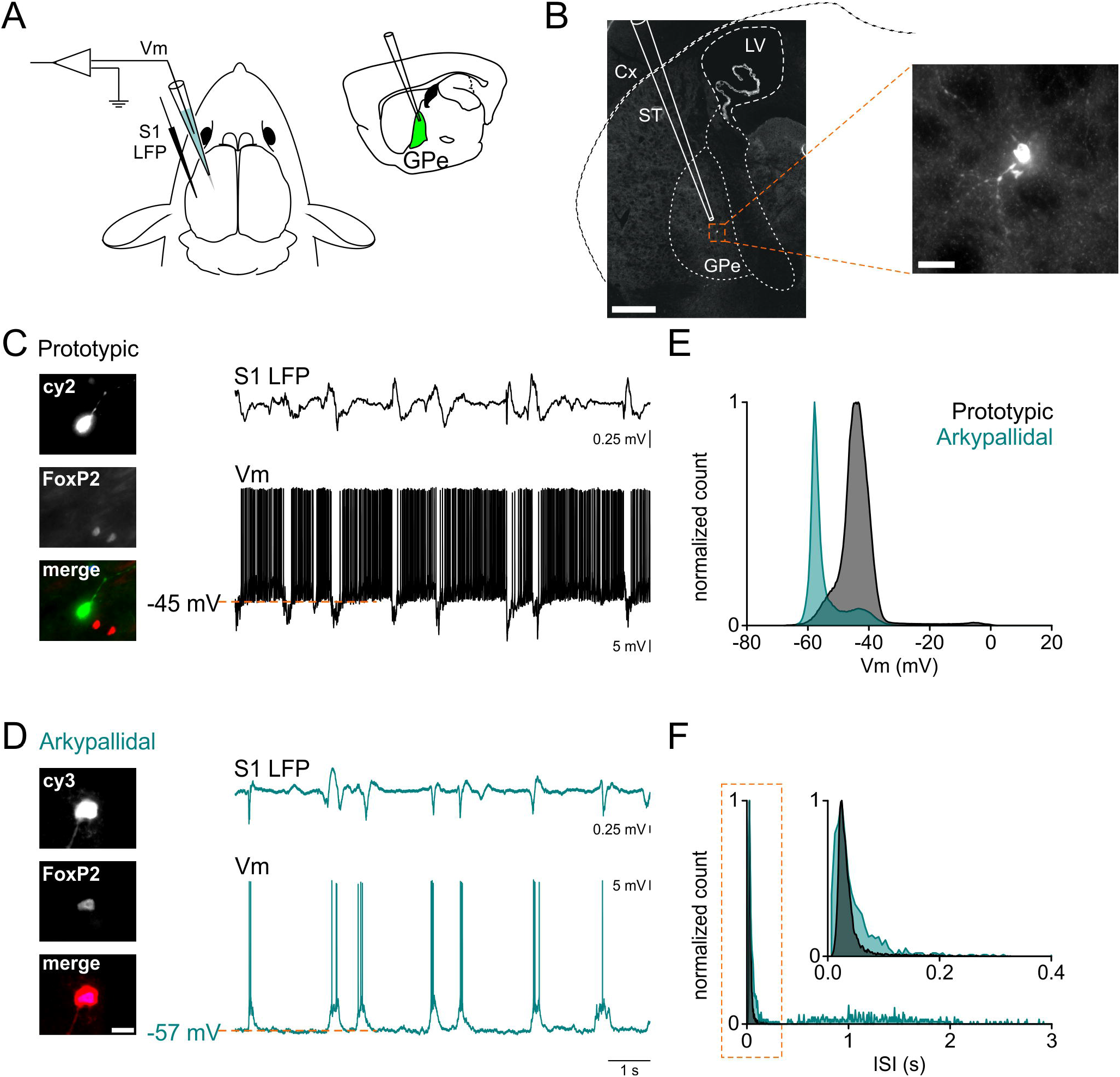
*In vivo* whole-cell recordings from prototypic and arkypallidal GPe cells. A. a scheme describing the experimental set up, including LFP cortical recordings (S1 LFP) and GPe whole-cell recordings. B. example of a GPe neuron labeled with biocytin following whole-cell recording (scale bar 500 μm). The right panel shows the recovered cell in higher magnification (scale bar 25 μm). C, *left panel*: identification of prototypic cells according to their expression of FoxP2: prototypic cells are negative to FoxP2. Scale bar images 25 μm (as indicated in D). C, *right panel*: Prototypic cells (black, top) fire at high frequencies and slow or pause their firing when the cortex (S1 LFP, top trace) is engaged in an up-state. D, *left panel*: identification of arkypallidal cells according to their expression of FoxP2: arkypallidal cells are positive to FoxP2. Scale bar images 25 μm. D, *right panel*: arkypallidal cells (turquoise) typically depolarize during cortical up-states. E. membrane potential histogram of a prototypic and an arkypallidal cell (same as in C and D). F. inter-spike interval (ISI) histogram for the cells presented in C-D. Prototypic cells in black, arkypallidal cells in turquoise, Cx cortex, ST striatum, GPe external globus pallidus, LV lateral ventricle, Vm membrane potential, LFP local field potential, S1 primary somatosensory cortex.

### Electrophysiological properties of GPe cells in vivo

We used whole-cell recordings *in vivo* to extract and compare the membrane properties of molecularly identified prototypic and arkypallidal cells. Prototypic cells (P) were significantly more depolarized than arkypallidal cells (A) (P: −46.09 ± 0.55, A: −60.60 ± 0.95 mV, p < 0.001), fired spontaneously at higher average rates (P: 14.20 ± 1.14, A: 0.58 ± 0.26 Hz, p < 0.001), and had larger sag ratios (P: 1.10 ± 0.01, A: 1.06 ± 0.01, p < 0.001) (n = 44 prototypic, 32 arkypallidal cells, Figure 2A-B). Other membrane properties such as input resistance (P: 0.22 ± 0.01, A: 0.23 ± 0.01 GΩ, p = 0.55) were not different between the groups. In order to identify cell types in an unbiased manner based on their electrophysiological properties, we performed unsupervised clustering of 259 recorded GPe neurons, as previously described for slice recordings (Abrahao and Lovinger, 2018) (Figure 2C). The clustered data showed an initial division into two main groups that corresponded to the molecularly defined subgroups of prototypic and arkypallidal cells (Figure 2C). The two groups differed in various electrical properties, including both subthreshold properties and features of their action potential (AP) firing (Figure 2D). Prototypic cells fired more regularly (lower CV_isi_) than arkypallidal cells (P: 0.99 ± 0.04, A: 1.31 ± 0.06, p < 0.001) and had smaller AP amplitudes (P: 47.30 ± 0.96, A: 56.29 ± 1.59 mV, p < 0.001). These properties accompanied the significant differences observed in their membrane potential (P: −46.14 ± 0.25, A: −58.34 ± 0.55 mV, p < 0.001), average spontaneous firing frequency (P: 14.22 ± 0.58, A: 1.08 ± 0.18 Hz, p < 0.001) and sag ratio (P: 1.12 ± 0.01, A: 1.05 ± 0.01, p < 0.001), already described for identified cells (n = 175 prototypic cells, n = 84 arkypallidal cells, Figure 2D). Input resistance (P: 0.23 ± 0.01, A: 0.22 ± 0.01 GΩ), AP half width (P: 0.60 ± 0.02, A: 0.66 ± 0.04 ms) and AP afterhyperpolarization (P: −5.59 ± 0.17, A: −6.22 ± 0.35 mV) did not differ between the classified groups (p > 0.153).

**Figure 2:**
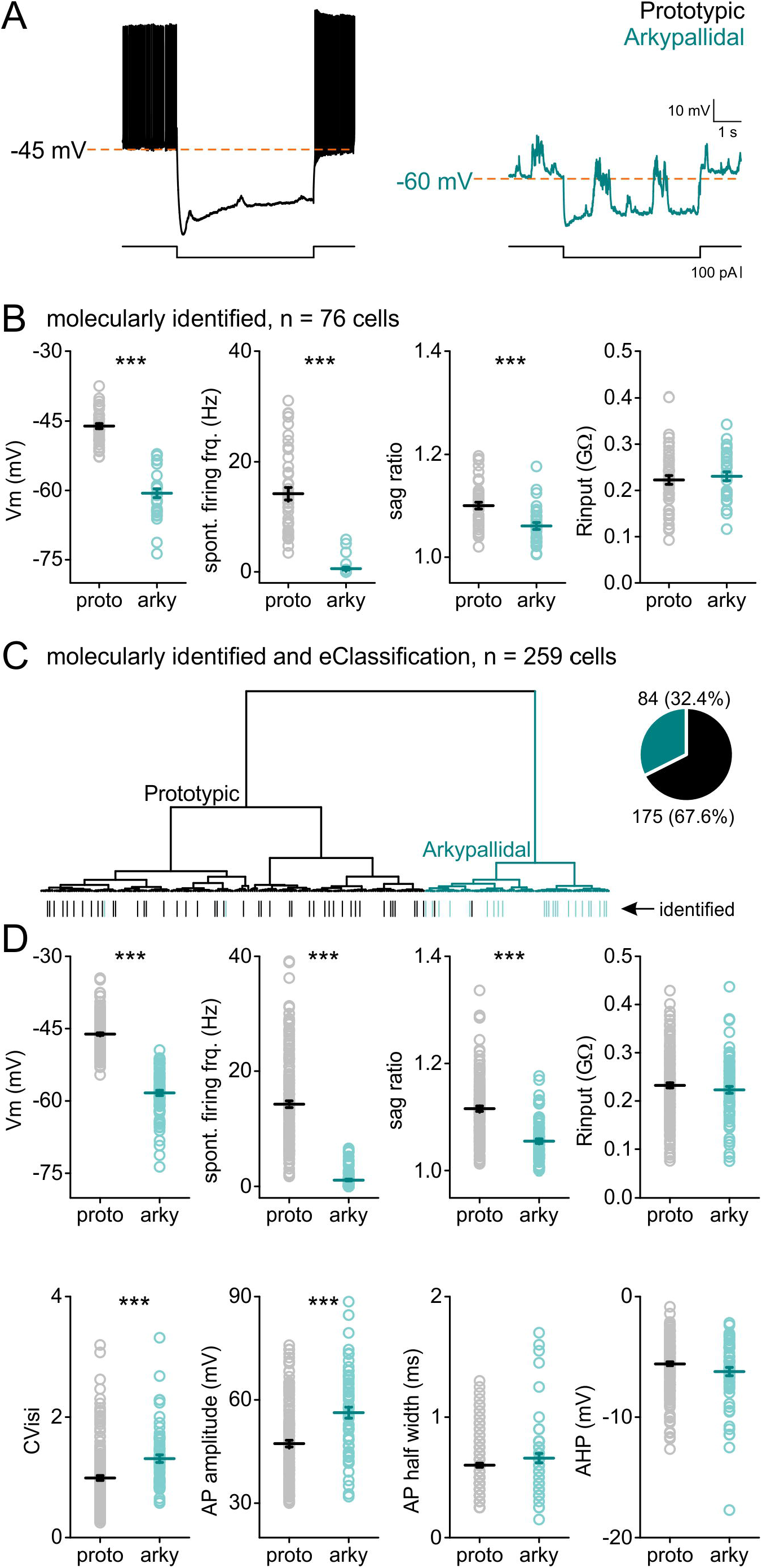
Membrane properties of identified and classified GPe neurons. A. examples of traces from prototypic (black, left) and arkypallidal (turquoise, right) cells showing the baseline activity for each cell, and responses to negative step current injection, exposing the sag in both populations. B. intrinsic properties of GPe cells identified either by immunostaining or by the “optopatcher” (n = 44 prototypic, n = 32 arkypallidal). C. dendrogram showing the classification of 259 recorded GPe cells. Inset: pie chart representing the distribution of recorded neurons. Note the marking of the identified cells at the bottom of the dendrogram. D. intrinsic properties of the GPe cells according to their classification by membrane potential, spontaneous firing frequency, and sag ratio. Prototypic cells in black, arkypallidal cells in turquoise, *** p<0.001. Data are presented as mean ± SEM. Statistical test: Mann whitney / two tailed T-test.

The hierarchical cluster analysis showed that while the classified prototypic and arkypallidal cells diverge very early on, prototypic cells can be further divided into two groups. Although no molecular markers were used to distinguish between the subgroups of prototypic cells, we compared the membrane properties of the three groups created by the classifier (Figure S2). The same electrophysiological properties used to separate prototypic and arkypallidal cells were used to further classify the two types of prototypic cells (P1 and P2). Both membrane potential (P1: −47.56 ± 0.28, P2: −44.45 ± 0.36, A: −58.34 ± 0.55 mV) and spontaneous firing frequency (P1: 8.73 ± 0.33, P2: 20.75 ± 0.69, A: 1.08 ± 0.18 Hz) were different between all three groups (p < 0.001). The sag ratio, however (P1: 1.12 ± 0.01, P2: 1.11 ± 0.01, A: 1.05 ± 0.01), was not different between the two prototypic groups (p = 0.93) but did differ between either of the prototypic groups and arkypallidal cells (p < 0.001). We thus named the two subgroups of the prototypic cells: ‘slow’ (SP, previously P1) and ‘fast’ (FP, previously P2) reflecting the differences in firing rate and membrane potential (n = 80 FP cells, and n = 95 SP cells). A three group comparison revealed differences in the regularity of spontaneous firing, with the fast prototypic cells having lower CV_isi_ than the other types (FP: 0.86 ± 0.05, SP: 1.09 ± 0.06, A: 1.30 ± 0.06, p < 0.001), as well as smaller AP amplitudes (FP: 42.73 ± 1.20, SP: 50.69 ± 1.30, A: 56.29 ± 1.59 mV, p < 0.01). No differences were found in input resistance (FP: 0.23 ± 0.01, SP: 0.23 ± 0.01, A: 0.22 ± 0.01 GΩ, p = 0.58), AP half width (FP: 0.59 ± 0.02, SP: 0.61 ± 0.03, A: 0.67 ± 0.04 ms, p = 0.24), and AP afterhyperpolarization (FP: −5.33 ± 0.27, SP: −6.05 ± 0.26, A: −6.26 ± 0.36 mV, p = 0.11). These data show that by extracting membrane properties recorded *in vivo*, GPe neurons can be classified into the previously defined prototypic and arkypallidal cells. The same classification algorithm also indicates the existence of subgroups within the prototypic cell population with distinct electrophysiological properties.

### Both GPe cell types receive excitatory and inhibitory inputs during slow wave activity

The opposite modulation of GPe cell types during up-states (recorded in cortical LFP, Figure 1) could be due to differences in their membrane properties (Figure 2) as well as differences in the synaptic inputs they receive during up-states. In order to explore these two possibilities, we manipulated the membrane potentials of recorded neurons by injecting negative and positive holding currents via the patch-pipette (Figure S3). When hyperpolarizing prototypic cells, the activity coinciding with the up-states reversed polarity and was now depolarizing, occasionally enabling firing during cortical up-states. This indicates that during up-states prototypic cells do not only receive inhibitory input, but also an excitatory component. Conversely, depolarization of arkypallidal cells reversed their modulation during cortical up-states, which now resulted in hyperpolarization and reduction in firing rate, as observed in prototypic cells (Figures 3 and S3). These data suggest that during slow wave activity, both prototypic and arkypallidal cells receive a barrage of mixed excitatory and inhibitory inputs, and that the modulation of their spiking during up-states depends on their respective membrane potentials and baseline activity levels.

**Figure 3:**
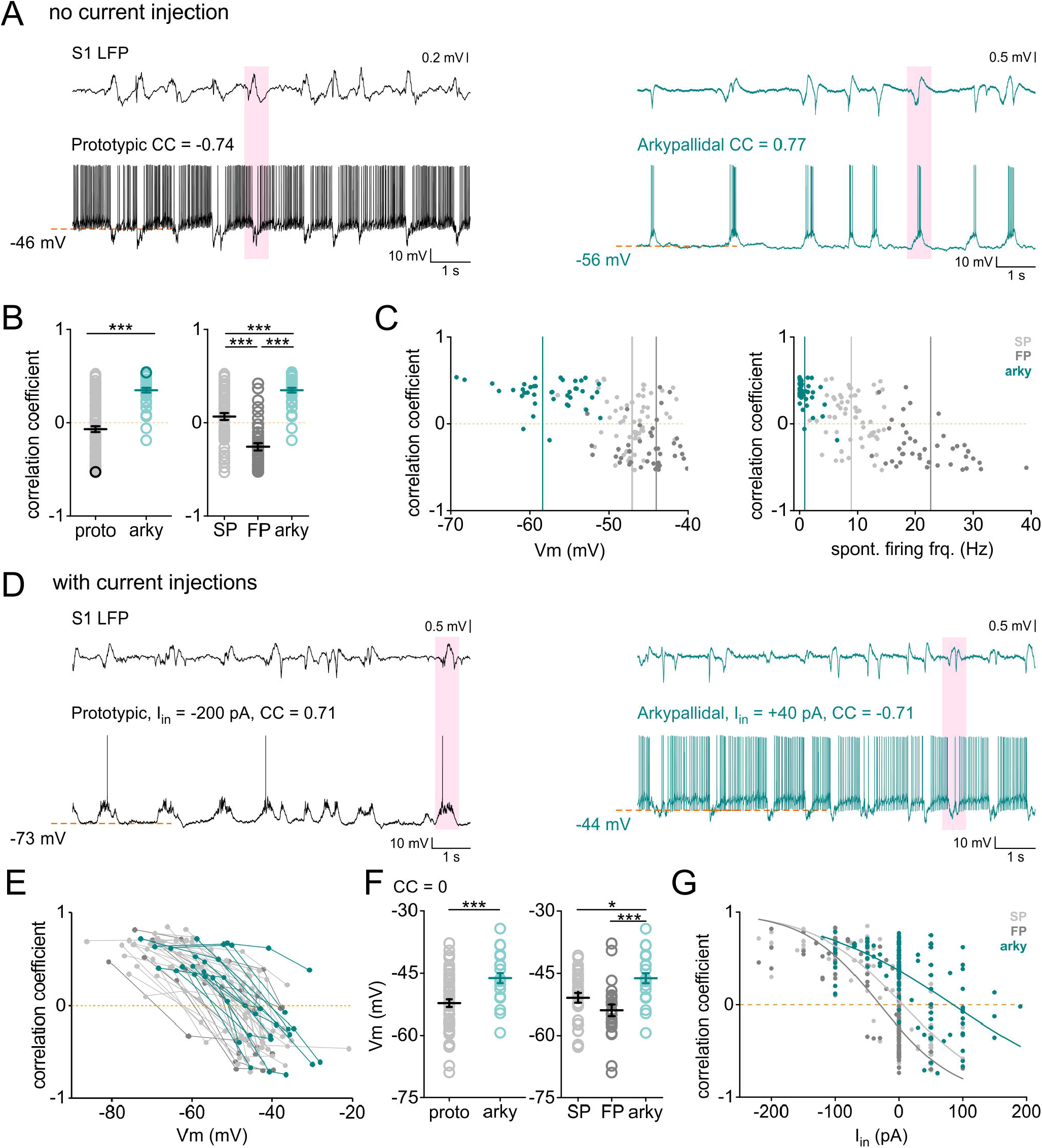
Modulation of membrane potential dynamics in GPe cells during slow wave activity. A. examples of traces recorded in prototypic (black) and arkypallidal cells (turquoise) without modulation of membrane potential by current injection (I_in_ = 0) and their correlation to the LFP recorded in S1. Note shaded area showing the cortical up-state and the corresponding activity in the whole-cell recorded GPe neuron. B. correlation coefficient of the cortical LFP and the cell activity filtered between 0.4-1.6 Hz, organized according to the classifier into 2 (left) or 3 (right) types of GPe neurons. Darker circles in the left panel corresponding to the examples in A. C. distribution of the correlation coefficient values according to membrane potential (left) and spontaneous firing frequency (right) of the cells organized by 3 groups. Lines in each graph indicate the mean value per group. D. examples of traces recorded in GPe cells injected with holding currents: hyperpolarized prototypic cell (left) and depolarized arkypallidal cell (right). Note the change in polarity of the correlation coefficient values for each cell type compared with A. The pink shaded areas show an example of the cortical up-state and the corresponding activity in the recorded neuron. E. correlation coefficients of all cells that were recorded in at least 3 different holding currents, organized into 3 cell-types. F. estimation of the membrane potential value in which the correlation coefficient shifts from positive to negative values, organized according to the classifier to 2 (left) or 3 (right) groups. G. distribution of all correlation coefficient values recorded in all protocols (with and without membrane potential modulation by current injections). The plot is superimposed by sigmoidal fitting between the maximum and minimum values of the correlation coefficient 1 and −1. CC: correlation coefficient, I_in_: current injected, Prototypic cells represented in black, arkypallidal cells in turquoise, slow prototypic (SP) in light grey, fast prototypic (FP) in dark grey. *** p<0.001, * p<0.05. Data are presented as mean ± SEM. Statistical test: Mann-Whitney / Kruskal-Wallis.

Although both cell types receive inhibitory and excitatory inputs during up-states, the respective magnitudes of these components could differ and shape the responses in a cell-type specific manner. In order to quantify such differences we calculated the correlation coefficient between the cortical LFP and the membrane potential of GPe cells (Figure 3, See Methods). This analysis enabled us to determine the polarity and degree of up-state modulation of whole-cell recorded neurons. The correlation coefficient of classified prototypic and arkypallidal cells differed in both magnitude and polarity (Figure 3A-C, P: −0.07 ± 0.03, A: 0.35 ± 0.03, p < 0.001). While almost all arkypallidal cells (96.7%) had positive correlation coefficients indicating depolarization during up-states, prototypic cells were, on average, only weakly and negatively modulated by cortical activity (the average correlation coefficient was slightly negative). Correlation coefficient values also differed when recorded neurons were further divided into three subpopulations (arkypallidal, slow prototypic, and fast prototypic cells). Most fast prototypic cells (89.6%) had negative correlation coefficient values with the cortical LFP, while slow prototypic cells had both positive (61%) and negative (39%) correlation coefficient values, which averaged around zero (SP: 0.07 ± 0.04, FP: −0.26 ± 0.04, A: 0.35 ± 0.03, p < 0.001, Figure 3B-C). This analysis further shows that the three GPe subpopulations responded differently to the barrages of synaptic input during slow wave oscillations.

In a subset of neurons, we recorded the ongoing activity of the same cells at different membrane potential values by injecting different holding currents via the patch-pipette (Figure 3D-E, S3), enabling us to calculate the correlation coefficient for each condition. We could then approximate the membrane potential at which correlation coefficient values changed from positive to negative, thereby indicating a “functional reversal potential” for the compound inputs during up-states. This value was more depolarized for arkypallidal cells compared to prototypic cells (P: −52.24 ± 0.93, A: −46.17 ± 1.21 mV, p < 0.001 for two group comparison, n = 55 prototypic, n = 23 arkypallidal cells). Arkypallidal cells differed from both prototypic subgroups (SP: −50.92 ± 1.19, FP: −53.95 ± 1.42, A: −46.17 ± 1.21 mV, p < 0.02) and no difference was found between the two prototypic subtypes (p = 0.21, n = 31 FP, n = 24 SP, Figure 3F). Most recorded neurons, regardless of their type, were found to be positively correlated with the cortical LFP during negative current injections, and vice versa, negatively correlated with the cortical LFP during positive current injections (Figure 3G).

Put together, these results suggest that prototypic and arkypallidal cells receive different compositions of excitatory and inhibitory synaptic inputs during slow wave activity. The modulation of their respective activities is, therefore, determined both by cell-type specific membrane properties and synaptic inputs. We next aimed to dissect the synaptic inputs to the respective GPe cell types from the local GPe circuitry, the STN, and the two types of striatal MSNs.

### Prototypic cells inhibit arkypallidal cells

Intra-pallidal connectivity may play an important role in GPe function, especially due to the perisomatic location of the local inhibitory synapses (Gross et al., 2011). Anatomical data (Sadek et al., 2007) and *ex vivo* recordings in slices (Bugaysen et al., 2013) have revealed only sparse connectivity amongst GPe cells, but it has been predicted that a primary intra-pallidal synaptic pathway exists from prototypic to arkypallidal cells (Nevado-Holgado et al., 2014). To study the synaptic interactions between GPe cells, we virally expressed ChR2 in the GPe of NKX2.1-Cre mice (Figure 4A-B). During *in vivo* recordings, we activated neurons with an optic fiber placed dorsally to the GPe (Figure 4A-B). ChR2-expressing prototypic cells responded to photostimulation with strong, reliable and sustained depolarization, superimposed by APs, even with very low light intensities (less than 0.2 mW, Figure 4D, G). The onset of light responses indicated a direct optogenetic excitation of recorded prototypic cells (0.43 ± 0.08 ms, n = 10, Figure 4 H). In contrast, arkypallidal cells were strongly inhibited by photostimulation (Figure 4 F, G and I) with onset delays suggesting monosynaptic inhibition (4.71 ± 0.37 ms, n = 5, Figure 4 H). The reversal potential of inhibitory responses was ~−75 mV, corresponding to the expected value for GABAergic inhibition (Figure S4). In two cases we recorded from a prototypic cells (FoxP2 negative) that did not virally express ChR2 (Figure 4E, G and J). Photostimulation of neighboring prototypic cells induced inhibitory synaptic responses in these cells (Figure 4 E, G and J, onset latency similar to arkypallidal cells, Figure 4 H), showing the existence of inhibition among prototypical cells. These results show that prototypic cells provide strong and reliable inhibition to arkypallidal cells and are also interconnected among themselves. It also suggests that in addition to afferent excitation, arkypallidal cells are also disinhibited during up-states due to the reduction in spiking of prototypic cells.

**Figure 4:**
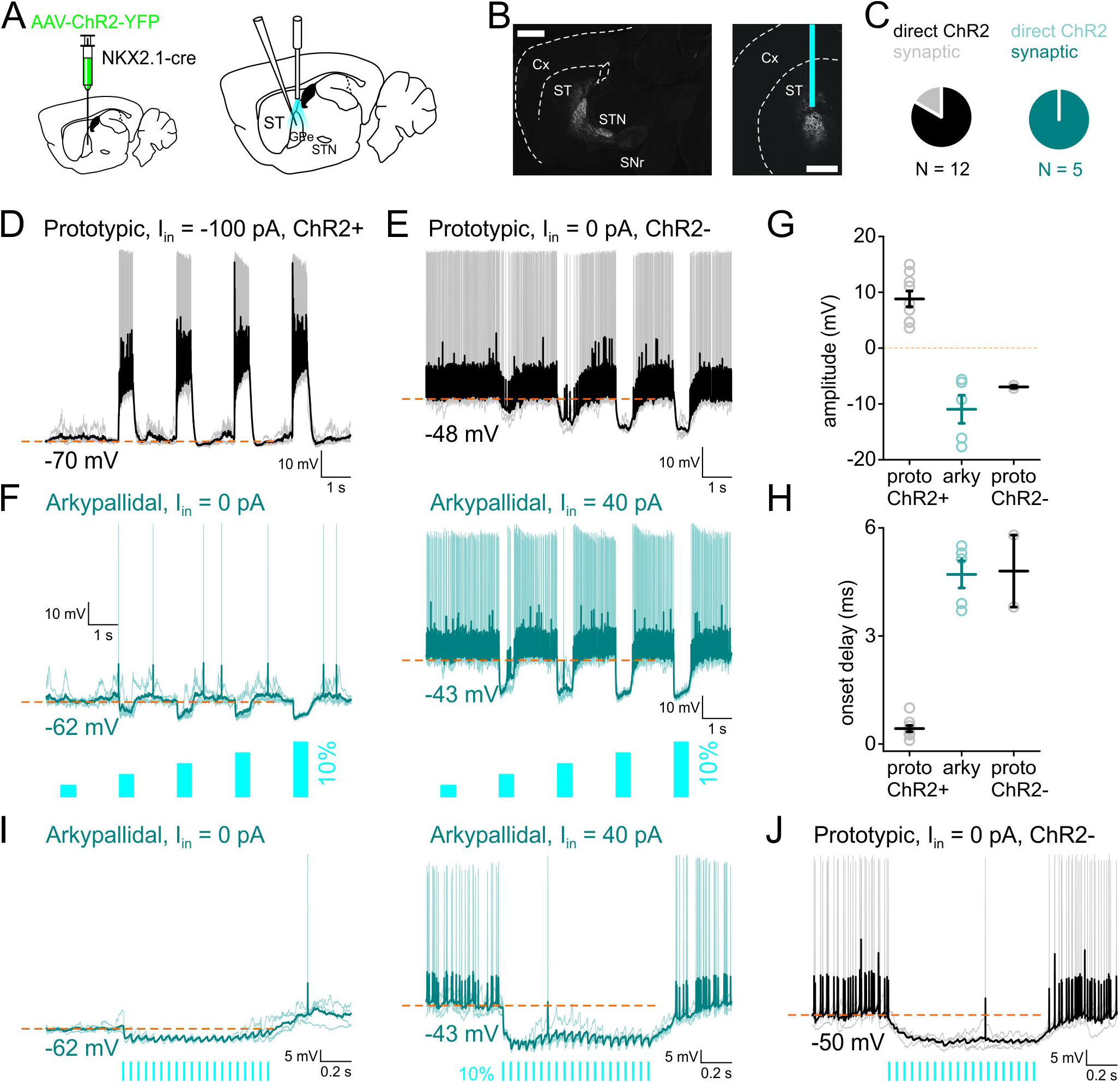
Lateral inhibition within the GP. A. a scheme of the experimental set up: viral injections in GPe (left), photostimulation and whole-cell recording (right). B. sagittal (left, scale bar 1 mm) and coronal (right, scale bar 1 mm) sections showing the virus expression and fiber location. C. pie charts representing the distribution of prototypic (black) and arkypallidal (turquoise) cells responding directly or indirectly (synaptically) to photostimulation. D. example of traces recorded in prototypic cells (black) expressing ChR2, held at −70 mV responding directly to photostimulation. E. example of traces recorded in a prototypic cell (black) which did not express ChR2, synaptically inhibited following photostimulation. For D-E, light stimulation is as indicated in F. F. example of traces recorded in arkypallidal cells (turquoise) at rest (left) or depolarized (right) during photostimulation. For both D-F traces are presented as raw traces in faint color overlaid with the average trace in darker color. G. light response amplitude in prototypic and arkypallidal cells. H. onset delay in prototypic and arkypallidal cells. I-J. examples of traces recorded in arkypallidal cell at rest (I, left), depolarized with current injection (I, right) and prototypic cell (J) to photostimulation with a train (20 pulses of 10 ms at 20 Hz). Prototypic cells represented in black, arkypallidal cells in turquoise.

### STN provides afferent excitatory input to both GPe cell types

The STN is reciprocally connected to the GPe and has been considered to be the main source of excitatory input to the GPe (Kita et al., 1983; Kita and Kitai, 1991; Pamukcu et al., 2020). To study the STN inputs to GPe, we expressed ChR2 in STN neurons using retrograde viral transduction in vglut2-Cre mice (Figure 5 A, see methods). We then recorded the responses of GPe cells to photostimulation of STN cells through a fiber placed above the STN (Figure 5A-B). All recorded GPe cells responded to STN photostimulation (Figure 5 C), however, there were differences in responses properties. The amplitude of the initial response phase was not significantly different between prototypic and arkypallidal cells (P: 13.62 ± 1.43, A: 9.12 ± 1.59 mV, n = 19 prototypic and 6 arkypallidal cells, p = 0.055, Figure 5 D-F), nor was there a difference in onset delays (P: 4.75 ± 0.14, A: 4.35 ± 0.1 ms, p = 0.14, Figure 5 G). In contrast, photostimulation with a 500 ms light pulse induced a sustained depolarization in prototypic cells (Figure 5 D) but only a transient response in arkypallidal cells (Figure 5 E). The response amplitudes measured at the end of the light pulse were strongly reduced in arkypallidal cells compared with their initial response amplitudes (start: 9.12 ± 1.59, end: 3.49 ±1.02 mV, p < 0.001), but not in prototypic cells (start: 13.62 ± 1.43, end: 14.32 ± 1.59 mV, p = 0.61, Figure 5 F). Photostimulation of the STN while holding GPe cells at −45 mV resulted in continuous depolarization of prototypic cells (Figure 5 H), in contrast to a brief depolarization followed by sustained hyperpolarization of arkypallidal cells (Figure 5 I), likely to originate from activated prototypic cells (Figure 4). Nevertheless, photostimulation of STN with a high frequency train (20 pulses at 20 Hz, Figure S5) induced depolarizing responses that enabled firing of action potentials in both cell types. These results show that while STN provides excitatory synaptic input to both GPe populations, the impact of this input is cell-type specific and shaped by intra-pallidal connectivity.

**Figure 5:**
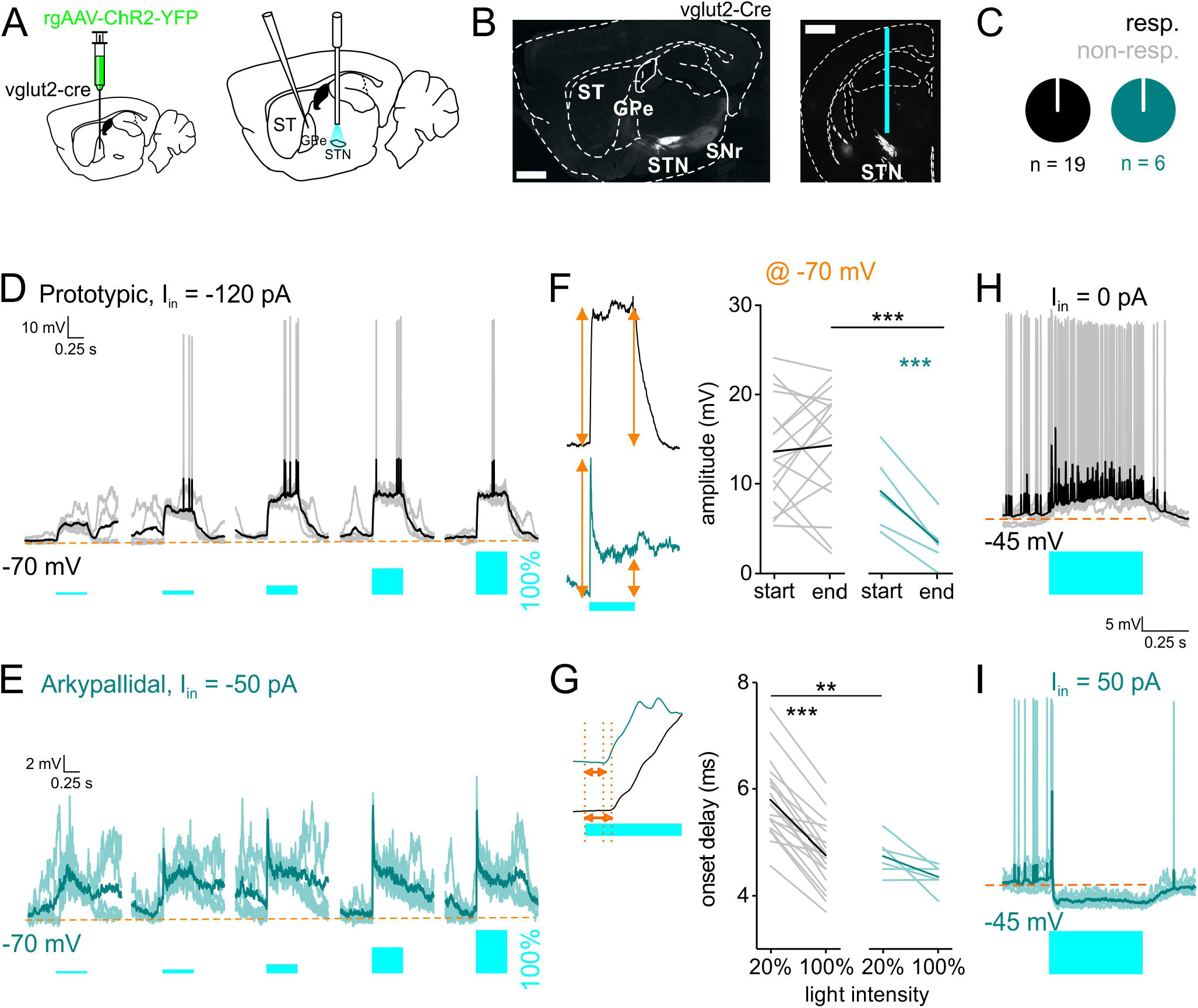
STN input to both GPe cell types. A. a scheme of the experimental set up, retrograde viral injections in GPe (left), photostimulation in STN and whole-cell recording in GPe (right). B. sagittal (left, scale bar 1 mm) and coronal (right, scale bar 1 mm) sections showing the specific viral expression in STN. C. pie charts showing that all recorded GPe cells responded to STN photostimulation. D-E. Responses of prototypic (D) and arkypallidal (E) cells to STN photostimulation with 500 ms light pulses of increasing light intensity (4, 8, 20, 60 and 100%). Recorded neurons were held at −70 mV by injection of negative holding currents (I_in_). F. quantification of the amplitude, at the beginning and end of the light step, at maximal light intensity. Inset showing an example of the measured parameter (orange). G. quantification of the onset delay to light stimulation. Inset showing an example of the measured parameter (orange). H-I. responses to 500 ms light activation when prototypic cells (H) and arkypallidal cells (I) are held at −45 mV. Note the initial spiking with light onset in arkypallidal cells. Prototypic cells in black, arkypallidal cells in turquoise. ** p<0.01, *** p<0.001. ST striatum, GPe external globus pallidus, SNr substantia nigra pars reticulata, STN subthalamic nucleus. Data are presented as mean ± SEM. Statistical tests: Paired sample T test / Two sample T test.

### Target selectivity in striato-pallidal inhibition

The major source of inhibition to the GPe is attributed to the striatum and particularly to indirect pathway MSNs (iMSNs), however, axon collaterals of direct pathway MSNs (dMSNs) were also shown to project to the GPe (Cazorla et al., 2014; Kawaguchi et al., 1990; Wu et al., 2000). In order to understand how the two types of MSN inhibit GPe subpopulations, we expressed ChR2 in either direct or indirect pathway MSNs and recorded the synaptic responses in GPe neurons to striatal photostimulation. To study the inputs of iMSNs to the GPe, we expressed ChR2 in iMSNs either virally using D2-Cre or A2A-Cre mice (Figure 6 A-B), or in D2-Cre mice crossed with a ChR2 reporter mouse (Ai32, Figure S6). As expected, in all experimental groups, prototypic cells were strongly inhibited by photostimulation of iMSNs (Figure 6 C-E). It was thus expected that inhibition of prototypic cells may cause depolarization of arkypallidal cells via disinhibition (Figure 4), however, in all cases (Figure 6 C), arkypallidal cells responded to photostimulation by an initial hyperpolarization (Figure 6 F-I), indicating direct inhibition by iMSNs. The initial hyperpolarization was often followed by a delayed depolarization (Figure 6 H), likely to originate from the reduced firing of neighboring prototypic cells (Figure 6 D). The amplitude of inhibitory responses (Figure 6 J) was larger in prototypic cells compared to arkypallidal cells held at similar membrane potential using current injections (P: −19.12 ± 0.54, A: −8.90 ± 1.37 mV, p < 0.001), and the onset delay (Figure 6 K) was not different (P: 7.34 ± 0.35, A: 8.60 ± 0.43 ms, p = 0.051, cells recorded in A2a-Cre or D2-Cre, n = 27 prototypic, n = 11 arkypallidal). Similar results were obtained in D2-ChR2 mice using the same experimental configuration (Figure S6). These data suggest that while both prototypic and arkypallidal cells receive inhibition from iMSNs, this inhibition is biased towards prototypic cells and can induce a delayed disinhibition of arkypallidal cells.

**Figure 6:**
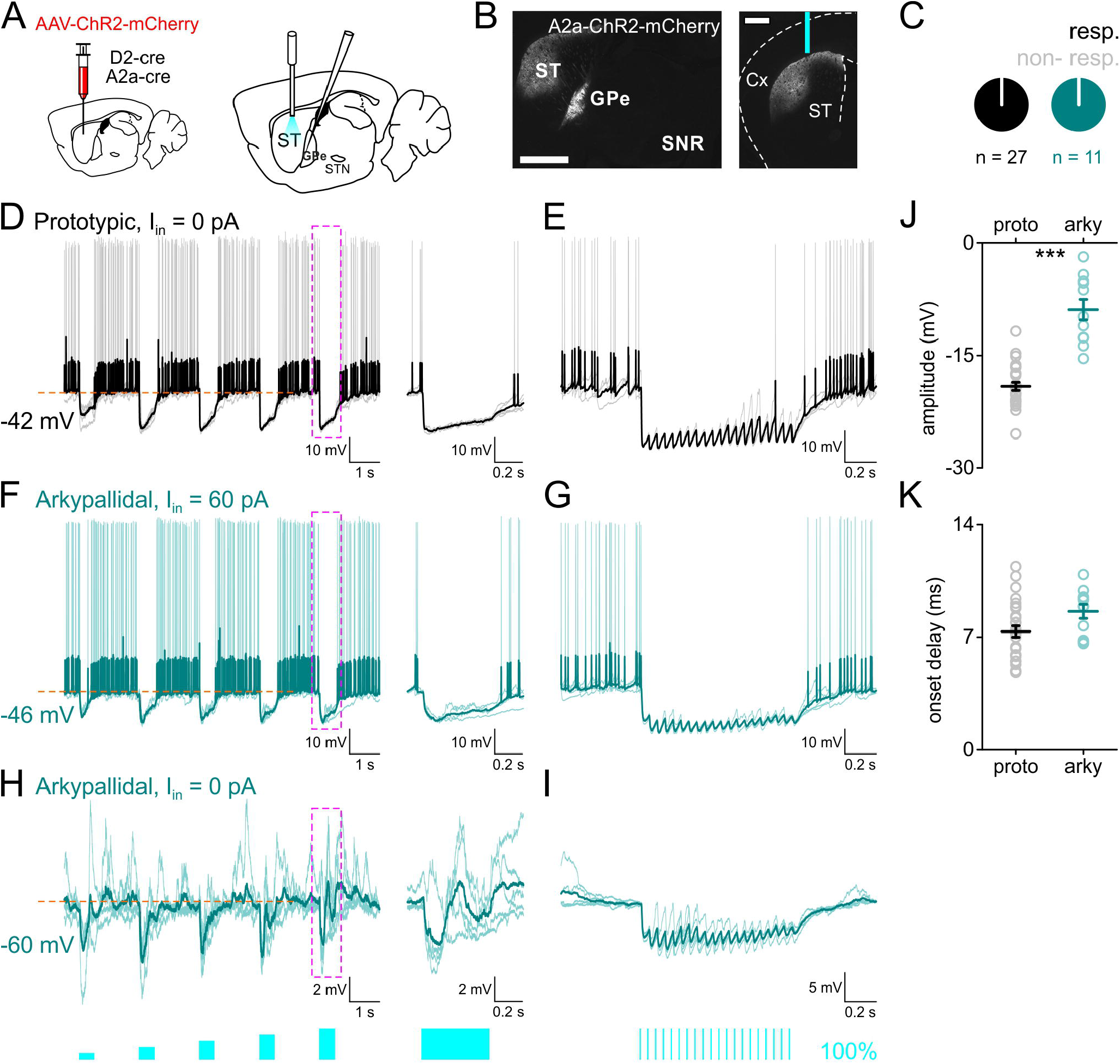
Indirect-pathway MSNs input to the GPe is biased towards prototypic cells. A. a scheme of the experimental set up: viral injections in striatum (left), photostimulation in striatum and whole-cell recording in GPe (right). B. sagittal (left, scale bar 1 mm) and coronal (right, scale bar 0.5 mm) sections showing the virus spread in the dorsal striatum and fiber location, and the typical projection of iMSNs to GPe. C. pie charts showing that all recorded GPe cells responded to iMSN photostimulation. D. responses of prototypic cells to 500 ms light stimulation of striatal iMSNs. Magenta dashed area expanded to the right of the trace. E. responses of prototypic cells to 20 Hz light stimulation of iMSNs. Baseline membrane potential is as indicated in D. F. same as in D but for depolarized arkypallidal cells. G. same as in E but for depolarized arkypallidal cells. H. same as in D but for arkypallidal cells at rest. I. same as in E but for arkypallidal cells at rest. J. light response amplitude in prototypic and arkypallidal cells held at similar membrane potentials. K. response onset delay to photostimulation in prototypic and arkypallidal cells held at similar membrane potentials. Prototypic cells in black, arkypallidal cells in turquoise. ST striatum, GPe external globus pallidus, SNr substantia nigra *pars reticulata*. Data are presented as mean ± SEM. Statistical tests: Two sample T test / Mann Whitney.

To study the inputs from dMSNs to GPe cells, we virally expressed ChR2 in the striatum of D1-Cre mice and positioned an optic fiber in the dorsolateral striatum (Figure 7 A-B). Out of 33 prototypic cells recorded, only 6 (18.18 %, 3 SP and 3 FP cells) showed measurable responses to photostimulation (Figure 7 C-E). In contrast, all arkypallidal cells were inhibited by dMSN photostimulation, as seen by short-latency hyperpolarization of membrane potential and suppression of action potentials (n = 5, Figure 7 C F-I). The amplitude of inhibitory responses was larger in arkypallidal than in prototypic cells (P: −0.58 ± 0.24, A: −10.99 ± 2.62 mV, p < 0.001, n = 33 prototypic and 5 arkypallidal cells, Figure 7 J), but no difference was found in onset delay (P: 8.6 ± 1.28, A: 8.08 ± 1.14 ms, p = 0.99, n = 6 prototypic and 5 arkypallidal cells, Figure 7 K). These results indicate a strong bias in dMSNs input to the GPe, with strong and prevalent inhibition of arkypallidal cells and only sparse and weak inhibition of prototypic cells.

**Figure 7:**
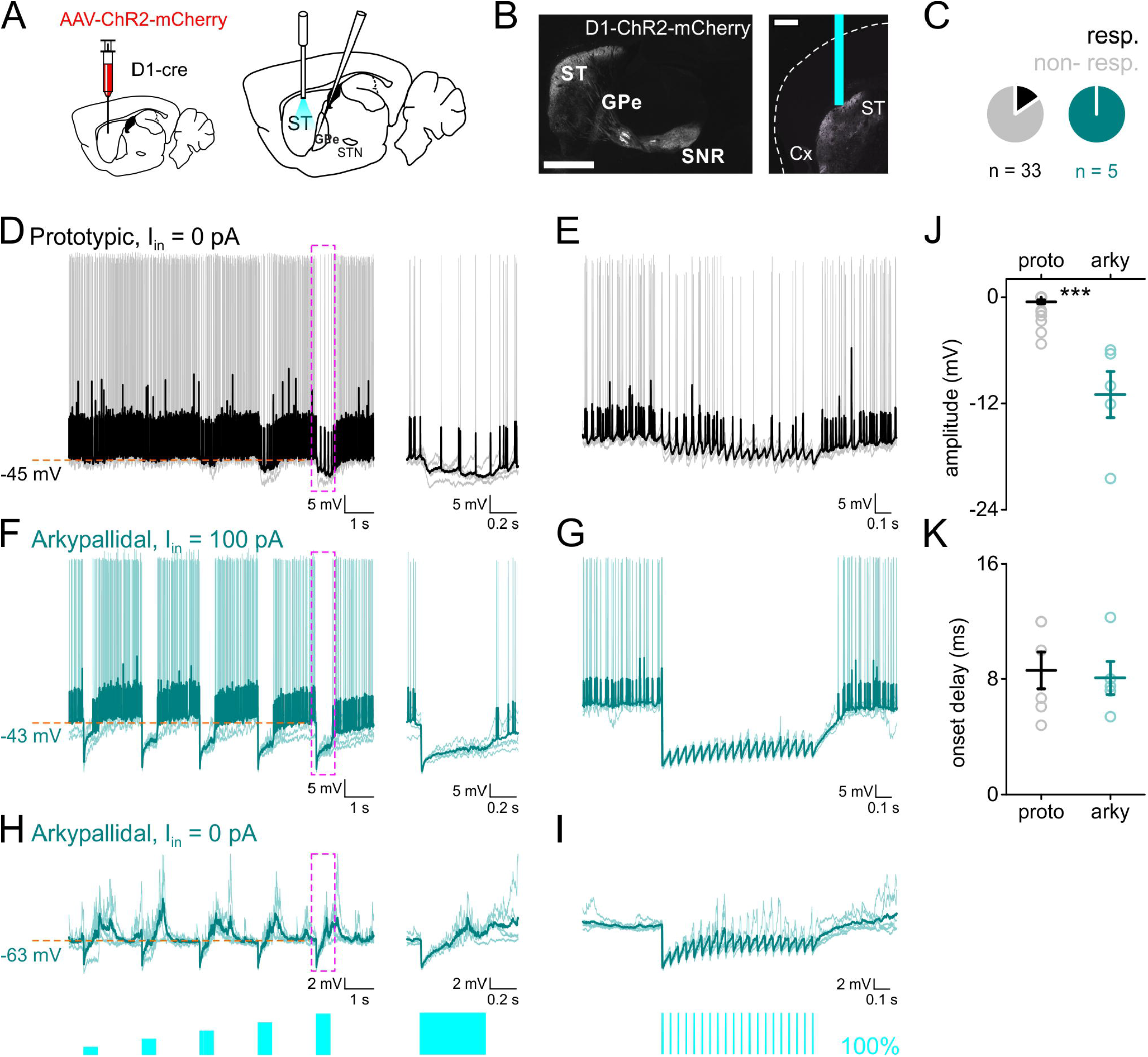
Direct-pathway MSNs target arkypallidal cells and almost avoid prototypic cells. A. a scheme of the experimental set up, viral injections in striatum (left), photostimulation in striatum and whole-cell recording in GPe (right). B. sagittal (left, scale bar 1 mm) and coronal (right, scale bar 0.5 mm) sections showing the viral transduction in dorsal striatum, the optic fiber location, and the typical projection of direct pathway MSNs to GPi and SNr. C. pie chart representation of the proportion of the cells responding to dMSN light activation. D. responses of prototypic cells to 500 ms light stimulation of striatal dMSNs. Magenta dashed area expanded to the right of the trace. E. responses of prototypic cells to 20 Hz light stimulation of dMSNs. Baseline membrane potential is as indicated in D. F. same as in D but for depolarized arkypallidal cells. G. same as in E but for depolarized arkypallidal cells. H. same as in D but for arkypallidal cells at rest. I. same as in E but for arkypallidal cells at rest. J. light response amplitude in prototypic and arkypallidal cells held at similar membrane potentials. K. response onset delay to photostimulation in prototypic and arkypallidal cells held at similar membrane potentials. Prototypic cells in black, arkypallidal cells in turquoise. ST striatum, GPe external globus pallidus, SNr substantia nigra *pars reticulata*. Data are presented as mean ± SEM. Statistical tests: Two sample T test / Mann-Whitney.

Our results show that the different GPe cell types receive input from both STN and striatum, however, there was pronounced target preference manifested in the response amplitudes, kinetics, and connection probabilities. Interestingly, arkypallidal cells receive reliable synaptic inputs from STN, dMSNs, and iMSNs, therefore linking them to the three main basal ganglia pathways (Figure 8).

**Figure 8:**
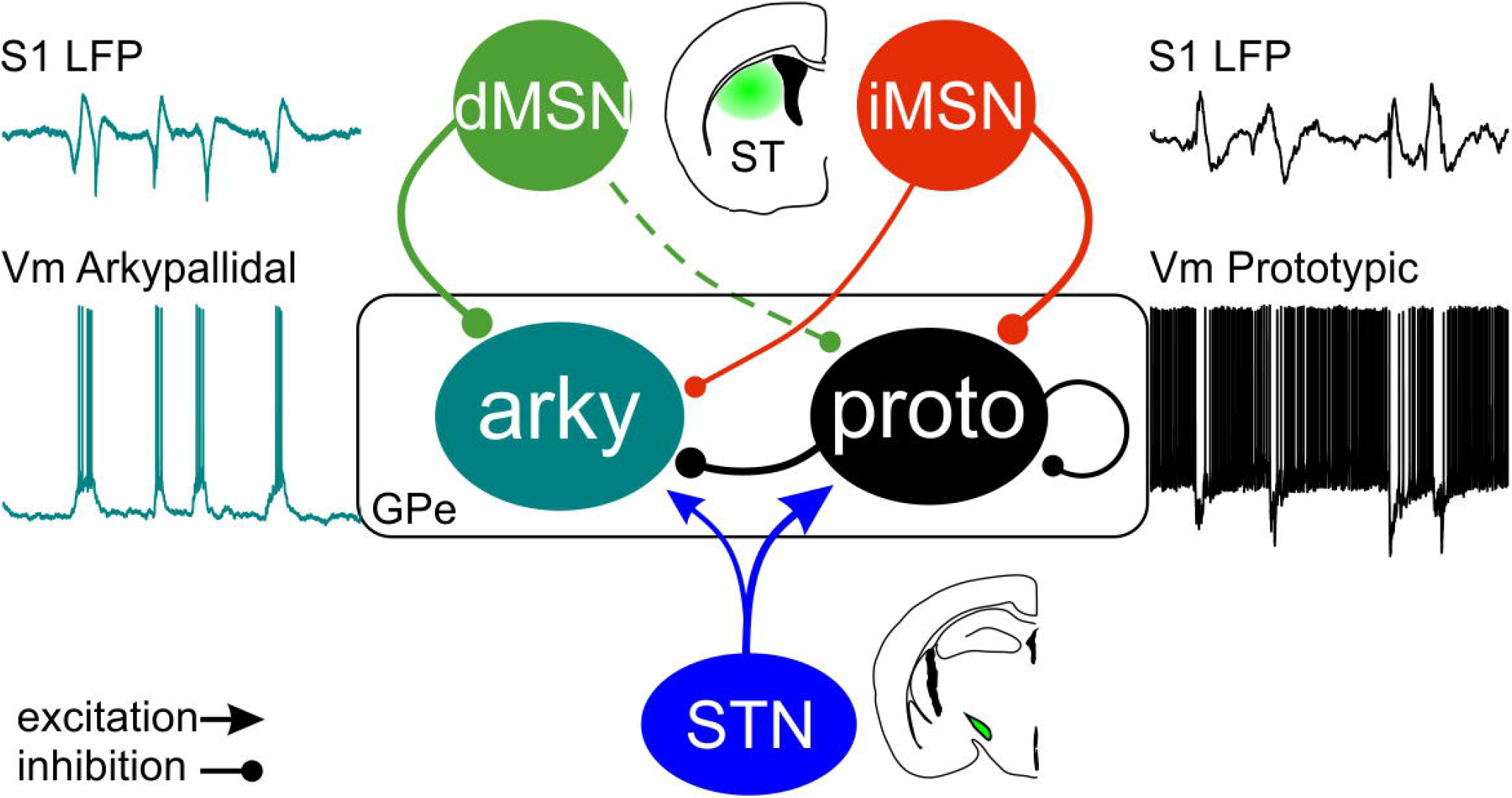
Summary scheme of synaptic inputs to GPe subpopulations. Prototypic cells (black) receive stronger input from iMSNs (red) and the STN (blue) and only sparse inhibition from dMSNs (green). Arkypallidal cells (turquoise) receive input from all basal ganglia pathways through direct and indirect pathway MSNs and the STN, as well as intra-pallidal inhibition from prototypic cells. S1 LFP: local field potential recorded in primary somatosensory cortex. Vm: membrane potential.

## Discussion

In this study, we used *in vivo* whole-cell patch clamp recordings in mice to study the membrane properties and network connectivity of GPe cells. To our knowledge this is the first report of such recordings in the GPe, enabling the study of both sub- and suprathreshold activity in the intact brain. Combining whole-cell recordings with optogenetics, we characterized the membrane properties and afferent inputs to the different GPe subpopulations. We show that during slow wave activity both prototypic and arkypallidal cells receive barrages of excitatory and inhibitory inputs that, together with their electrophysiological properties, pattern their spontaneous activity. Both GPe subpopulations receive input from STN and striatum, but there is a clear target preference that biases these afferent inputs, in particular the very weak input to prototypic cells from dMSNs. Notably, arkypallidal cells receive direct pathway information from dMSNs, indirect pathway information from iMSNs and prototypic cells, and hyper-direct input from the STN, thus placing them as integrators of the three main BG pathways (Figure 8).

We identified cells as prototypic or arkypallidal cells primarily according to their expression of the molecular markers FoxP2 and NKX2.1 thus dividing our recorded cells to prototypic and arkypallidal cells, without further subdivision. FoxP2 is an accepted molecular marker for the arkypallidal cells and it is rarely co-expressed with NKX2.1 or PV (Abdi et al., 2015; Dodson et al., 2015; Hernandez et al., 2015). The activity of molecularly identified GPe cells acquired using whole-cell *in vivo* recordings was in accordance with the activity described thus far using extracellular recordings (Abdi et al., 2015; Dodson et al., 2015; Mallet et al., 2012). Noteworthy, in some cases arkypallidal cells were completely silent under control conditions, as previously reported (Mallet et al., 2012). Using data obtained from molecularly identified cells we further classified cells based on their electrophysiological properties. Employing unsupervised clustering we could then compare the membrane properties for a total of 259 GPe cells and confirm the identity of the resulting groups according to the presence of the molecularly identified cells within them. Using this method, we explored the membrane properties of a larger number of GPe cells recorded *in vivo*. As expected, we found differences in spontaneous firing frequency and regularity of firing between our putative prototypic and arkypallidal cells, previously described with extracellular recordings (Abdi et al., 2015; Dodson et al., 2015; Mallet et al., 2012). In addition we found significant differences in membrane potential, sag ratio and action potential amplitude values, some of which were described using *ex vivo* recordings (Abdi et al., 2015; Abrahao and Lovinger, 2018; Hernandez et al., 2015; Mastro et al., 2014). The prototypic population could be further subdivided into two main groups which we named “slow” and “fast” prototypic cells based on their average spontaneous spiking frequency. Such division of the prototypic population was suggested in previous studies based on their differential expression of molecular markers and axonal projections (Abdi et al., 2015; Abecassis et al., 2020; Dodson et al., 2015; Hernandez et al., 2015; Mastro et al., 2014). In this study, however, we did not find a clear match between the prototypic subgroups to specific molecular markers, but this is an important topic for future studies.

By performing whole-cell recordings from GPe cells we could use current injections to study their afferent inputs during slow wave oscillations. Specifically, we evaluated the respective contribution of excitatory and inhibitory inputs to the GPe subpopulations during cortical up-states. We used the correlation coefficient between cortical slow wave activity and GPe cells membrane potential as an approximation of the compound excitatory-inhibitory ratio of the synaptic barrages. Although a more accurate estimation of this ratio could be obtained using voltage-clamp with QX314 and cesium in the pipette solution, such conditions would prevent extraction of the electrophysiological properties of recorded cells. Using the correlation coefficient analysis, we could show that both cell populations received mixed excitatory and inhibitory inputs during slow wave activity and that the excitation-inhibition balance was cell-type specific.

Our data show that prototypic cells provide strong and reliable inhibition of arkypallidal cells (Figure 4), implying that they are under tonic inhibition which is relieved transiently when prototypic cells are inhibited. We also showed that prototypic cells are interconnected by inhibitory synapses (Figure 4), however, the organization of this recurrent inhibition is yet unclear and will be subject to future studies. The afferent pathways to the GPe have been described previously (Albin et al., 1989; Cazorla et al., 2014; DeLong, 1990; Kawaguchi et al., 1990; Kita et al., 1983; Robledo and Feger, 1990; Wu et al., 2000), however, their impact on the different GPe subpopulations was unknown. We showed that STN provides excitatory input to both prototypic and arkypallidal cells, however, its impact on the activity of the respective subpopulations was cell-type specific. Excitation of arkypallidal cells was curtailed by strong inhibition from prototypic cells, resulting in only a brief response at the onset of STN activation. The differential responses to STN photostimulation may also reflect differences in other synaptic properties of STN input to the GPe subpopulations.

Striatal input was also biased with respect to the postsynaptic GPe cell types. As expected, iMSNs strongly inhibited prototypic cells, however, they also provided reliable, although weaker, inhibition to arkypallidal cells. Surprisingly, we found that dMSNs target only a small fraction of prototypic cells and with much weaker inhibition compared with arkypallidal cells (figure 7). This target selectivity in the GABAergic inhibition from dMSNs to arkypallidal cells is intriguing, in face of the selective excitation of prototypic cells by Substance P (Mizutani et al., 2017). Other inputs to the GPe such as from cortical regions (Abecassis et al., 2019; Karube et al., 2019) were not explored in this paper, but are also likely to shape GPe activity.

Prototypic cells project downstream and inhibit the BG output structures but also other structures such as the STN (Bevan et al., 2002; DeLong, 1990; Smith et al., 1998), striatum (Bevan et al., 1998; Mallet et al., 2012; Mastro et al., 2014; Saunders et al., 2016), thalamic nuclei (Hazrati and Parent, 1991; Mastro et al., 2014) and the substantia nigra *pars compacta* (Mastro et al., 2014; Paladini et al., 1999). Within the GPe they exert local inhibition onto arkypallidal cells as well as themselves (Figure 4). Our findings show that prototypic cells are indeed targeted by iMSNs (Albin et al., 1989; Loopuijt and van der Kooy, 1985) and the STN (Kita et al., 1983; Robledo and Feger, 1990) as shown previously. STN activation results in reliable excitation of prototypic cells that, in turn, would inhibit downstream BG targets. iMSNs activation strongly inhibits the prototypic cells, thus disinhibiting the STN as well as neighboring arkypallidal cells. Our observations are, therefore, in line with the canonical role of prototypic GPe cells in the indirect pathway (Chiken et al., 2008; Nambu et al., 2000; Ozaki et al., 2017; Sano et al., 2013). The contribution of prototypic cell activation on structures outside the basal ganglia was not further explored in this paper.

The projection from GPe to striatum differs between prototypic and arkypallidal cells. Arkypallidal cells have been shown to target both striatal interneurons and MSNs (Mallet et al., 2012) while prototypic cells mainly target interneurons (Bevan et al., 1998; Saunders et al., 2016). Fractions of both GPe subtypes have been shown to express Npas1 (Abrahao and Lovinger, 2018; Hernandez et al., 2015), which labels GPe neurons that target both interneurons and MSNs (Glajch et al., 2016). Activation of STN provided strong and sustained excitation of prototypic cells, which would then inhibit striatal fast spiking interneurons (Bevan et al., 1998; Saunders et al., 2016), thus disinhibiting MSNs. In contrast, arkypallidal cells were briefly excited by STN activation, which would imply only a short window of inhibition onto MSNs. While prototypic cells constitute the majority of GPe neurons (Dodson et al., 2015; Mallet et al., 2012), the area covered by arkypallidal axons in the striatum is considerably larger than that of prototypic cells (Mallet et al., 2012). This suggests that arkypallidal cells have a strong and immediate impact on MSN activity, thus they may act as “stop cells” (Mallet et al., 2016). The impact of STN on the activity of MSNs via GPe pallidostriatal projections is, therefore, complex both spatially and temporally, requiring further investigation of the detailed functional organization. It was recently shown that arkypallidal cells receive direct input from motor cortex (Karube et al., 2019), suggesting that cortical activation may exert a dual impact on MSNs, by monosynaptic excitation followed by polysynaptic inhibition via GPe arkypallidal cells. Striatal inputs to the respective GPe populations showed a high degree of target preference, with iMSNs strongly inhibiting prototypic, but also arkypallidal cells. Interestingly, dMSN collaterals in GPe selectively inhibited arkypallidal cells, thereby suppressing the “stop signal” provided by this population. These data show that both direct and indirect striatal pathways shape GPe activity but they do so via different pathways in distinct manner.

In this paper we describe the electrical properties and synaptic organization of two main GPe neuronal populations, revealing cell-type specific afferent inputs. We show input bias to the GPe subpopulations which supports distinct functional roles. In future studies it will be important to elucidate the functional contribution of these pathways in sensorimotor function and their alteration leading to dysfunction.

## Acknowledgments

We thank Elin Dahlberg and Kristoffer Tenebro Berglund for technical help, Ole Kiehn, Gilberto Fisone, Konstantinos Meletis and Jens Hjerling-Leffler for mice. We also thank Sten Grillner, Abdel El Manira, and members of the Silberberg lab for comments and discussions. This work was supported by the Knut and Alice Wallenberg Foundation (KAW 2014.0051), the European Research Council (ERC 282012), the Swedish Brain Foundation (Hjärnfonden FO2018-0107), the Swedish Medical Research Council (VR-M 2015-02403), Karolinska Institutet Strategic Program for Neuroscience (StratNeuro), and grants from Karolinska Institutet.

## Author contribution

M.K. and G.S. conceived and planned the experiments. M.K. performed the experiments and analyzed the data. M.K. and G.S. wrote the manuscript.

## Materials and Methods

All experiments were performed according to the guidelines of the Stockholm municipal committee for animal experiments under an ethical permit to G.S. (N12/15). D1-Cre (EY217 line), D2-Cre (ER44 line), Adora2a-Cre (KG139 line, GENSAT), vglut2-Cre (Slc17a6^tm2(cre)Lowl^/J), PV-Cre (B6;129P2-Pvalbtm1(cre)Arbr/J) and NKX2.1-Cre mice (C57BL/6J-Tg(Nkx2-1-cre)2Sand/J, the Jackson laboratory) were used for virus injections. In some cases, Cre lines were crossed with the Channelrhodopsin (ChR2)-YFP reporter mouse line (Ai32, the Jackson laboratory) to induce expression of ChR2 in a specific cell population. Mice of both sexes were housed under a 12-hour light-dark cycle with food and water ad libitum. All experiments were carried out during the light phase.

### Virus injections

Mice, 6-8 weeks old, were anesthetized with isoflurane and placed in a stereotaxic frame (Stoelting). Craniotomy coordinates and injections volumes: **ST** (AP: 0.7, ML: 2.25, DV - 3.1) 0.5 - 1 μl of virus. **GPe** (AP: −0.35, ML: 2.15, DV: −3.65) 0.25 μl of virus. Viruses: AAV5.EF1.dflox.hChR2(H134R)-mCherry.WPRE.hGH or AAV5.EF1.dflox.hChR2(H134R)-eYFP.WPRE.hGH, rgAAV-EF1a-double floxed-hChR2(H134R)-EYFP-WPRE-HGHpA, addgene. Injections were done using a micropipette at 0.1 μl min^−1^ (Quintessential Stereotaxic Injector, Stoelting). The pipette was held in place for 5 min before being slowly retracted from the brain. Temgesic was applied after surgery (0.1 mg/Kg).

### In vivo recordings

Experiments were conducted as described previously (Ketzef et al., 2017; Reig and Silberberg, 2014), briefly, 2-3 months old mice, usually 3 weeks following virus injections, were anaesthetized by intraperitoneal (IP) injection of ketamine (75mg/kg) and medetomidine (1 mg/kg) diluted in 0.9% NaCl. To maintain mice under anesthesia, a third of the dose of Ketamine was injected intraperitonaelly approximately every 2 hours or in case the mouse showed response to pinching or changes in EcoG patterns. Mice were tracheotomized, placed in a stereotactic frame and received oxygen enriched air throughout the recording session. Core temperature was monitored with feedback-controlled heating pad (FHC) and was kept on 36.5±0.5 °C. The skull was exposed and 3 craniotomies were drilled (Osada success 40): for cortical LFP recordings, GPe intracellular recordings and for optic fiber placement. Sensory cortex craniotomy coordinates: 1.5 mm posterior to bregma, 3.25 mm lateral to midsagittal suture. A bipolar tungsten electrode with impedances of 1-2 MΩ was inserted 1 mm deep from the surface. Signals were amplified using a Differential AC Amplifier model 1700 (A-M Systems) and digitized at 20 KHz with CED and Spike 2 parallel to whole-cell recording. An optic fiber was inserted for activation of input population. A craniotomy was drilled and the fiber was inserted (AP, ML, DV in mm) to either ST (0.7, 2.25, −2), STN (−2, 1.5, −4.5) or above GPe (−0.35, 2.2, −2.75). For patch clamp recordings, the craniotomy was performed 0.3-0.5 posterior to, and 4.25 mm lateral to bregma, and the dura was removed. Patch pipettes were pulled with a Flaming/Brown micropipette puller P-1000 (Sutter Instruments). Pipettes (6-9 MOhm, borosilicate, Hilgenberg), back-filled with intracellular solution, were inserted with a ~1200 mbar positive pressure to a depth of about 2.8 mm from the surface, after which the pressure was reduced to 25-35 mbar. From that point the pipette was advanced in 1 μm steps in depth (32 degrees angle), in voltage clamp mode. When a cell was encountered, the pressure was removed to form a Gigaseal, followed by application of a ramp of increasing negative pressure till cell opening was evident. Recordings were performed in current clamp mode. Intracellular solution contained (in mM): 130 K-gluconate, 5 KCl, 10 HEPES, 4 Mg-ATP, 0.3 GTP, 10 Na_2_-phosphocreatine, and 0.2-0.3 % biocytin (pH=7.25, osmolarity~285 mOsm). The exposed brain was continuously covered by 0.9% NaCl to prevent drying. Signals were amplified using MultiClamp 700B amplifier (Molecular Devices) and digitized at 20 KHz with a CED acquisition board and Spike 2 software (Cambridge Electronic Design). The membrane potential of a cell was the peak value of the all point membrane potential histogram collected for over a minute. The spontaneous firing frequency was calculated for a similar period of time. Inter-spike interval coefficient of variance (CV_isi_) was calculated as the variance of the Inter-spike interval normalized to the average Inter-spike interval. Input resistance was calculated as the slope of the steady-state voltage responses to current injection (−100 pA to 0 pA in steps of 20 pA for 5 sec each). For each current injection, values during cortical up and down states were extracted separately. Sag was extracted from hyperpolarization protocol in which the cell was first not injected with any current, establishing baseline, followed by injection of −100 pA for 5 seconds. The sag was taken as the maximal drop in voltage at the beginning of the hyperpolarizing step. Sag ratio was the result of diving the sag by the steady-state voltage responses of the hyperpolarizing step. AP half width was calculated at half distance between membrane potential baseline and average AP peak value. Afterhyperpolarization (AHP) was measured between the membrane potential baseline to the average AP hyperpolarization peak value. For all these properties only recordings that were stable for a duration of at least a minute were taken; i.e. no drift in membrane potential or change in AP amplitude during the recording.

For classifying the intrinsic properties, we used data obtained from 259 GPe cells that included the molecularly identified cells. Unsupervised hierarchical cluster analysis was performed using Ward’s cluster method and Euclidean distance for parameters that were non-normally distributed. Since we could molecularly verify 2 clusters by the presence of the identified cells and the maximal drop in distance seen in the dendrogram divided the data into 2 groups, in most cases, the comparison is carried out between 2 groups.

Correlation coefficient was calculated from the filtered squared LFP signal recorded in S1 and the corresponding cell membrane potential in zero lag. Signals were filtered between 0.4–1.6 Hz using a second-order Butterworth filter (Abdi et al., 2015) before the correlation coefficient was extracted.

### Optogenetic stimulation

blue light (470 nm, mightex) was delivered though a cannula inserted to the craniotomy at these coordinates (AP, ML, DV in mm) to either ST (0.7, 2.25, −2.25), STN (−2, 1.5, −4.5) or above GPe (−0.35, 2, −2.75). Light activation protocols were triggered by spike2 program. Maximal LED light intensity at the tip of the cannula was 2 mW. All light stimulations are presented as percentages of the maximal light intensity. In subset of experiments we used the optopatcher (Katz et al., 2013; Ketzef et al., 2017) for identification of the cells recorded online.

### Cell labeling and immunohistology

During the recordings, the cells were loaded with biocytin (Sigma). At the end of the recording session, the mouse received an overdose of sodium pentobarbital (200 mg/kg I.P.), and transcardially perfused with 4% PFA in 0.01M phosphate buffer (PBS) pH 7.4. The brain was removed and kept for additional 2 hours in the fixative, after which it was transferred to 0.01 M PBS. The brain was transferred to and kept in 12% sucrose solution in 0.01M PBS overnight and < 20 μm cryo-sections were produced. sections were mounted on microscope gelatin coated slides and incubated for 2 hours in room temperature with cy2 / cy3 conjugated streptavidin (1:1000, Jackson ImmunoResearch Laboratories) in staining solution (1% BSA, 0.1% NaDeoxycholate and 0.3% triton in 0.01 PBS). After washes in PBS, slides were mounted on fluorescence microscope in order to locate the recorded cells. If a cell has been found, following the biocytin staining, we stained for FoxP2 (Rabbit anti mouse, abcam) expression (1:1000 in staining solution, overnight at 4 °C), followed by 2 hours incubation with secondary antibody (cy5 conjugated donkey anti rabbit, Jackson ImmunoResearch Laboratories, 1:500 in staining solution). Photomicrographs of the results were taken with Olympus XM10 (Olympus Sverige AB, Stockholm, Sweden) digital camera.

### Statistical analysis

Data are represented as mean ± SEM. The ns represent cells. Data distributions were first checked for normality (Shapiro-Wilk test) and analyzed accordingly. Normally distributed data were tested by one way ANOVA followed by post-hoc Tukey’s test analysis for multiple comparisons, and the unpaired and paired two-sample student’s T-test was used for two group comparisons. Non-normally distributed data were analyzed by Kruskal-Wallis test for multi-group comparisons followed by Mann-Whitney for two group comparison. Confidence level was set to 0.05. All statistical analyses were done in SPSS (IBM). Statistical test are reported in the figure legends. Experiments were not included if there was no virus expression, placement of fiber was not correct, or recordings were not stable, as described in the ‘*in vivo* recording’ section.

