## Supplementary material for "Differential Synaptic Input to External Globus Pallidus Neuronal Subpopulations *In Vivo*"

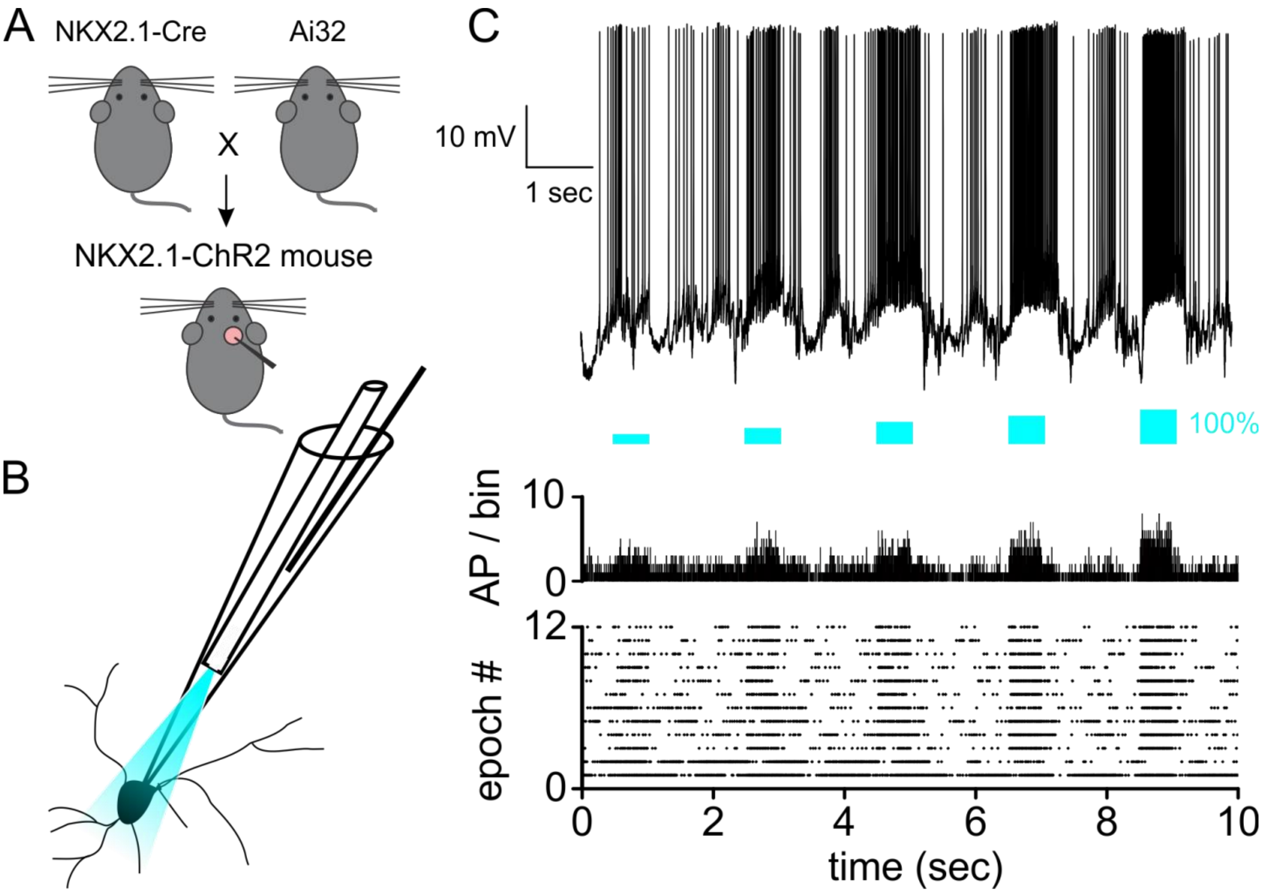


**Supplementary figure 1** (related to figures 1 and 2)**:** Online identification of prototypic cells using the optopatcher. A. crossing the NKX2.1-cre mouse with the Ai32 ChR2-YPF reporter mouse generated mice expressing ChR2-YPF in prototypic cells. B. a scheme of the optopatcher pipette tip which contains the optic fiber for photostimulation and the silver chloride wire for electrophysiological recording. C. example of whole-cell *in vivo* recording from a prototypic cell expressing ChR2 (top), photostimulation intensity profile (middle) and raster plot (bottom) recorded using the *optopatcher*.

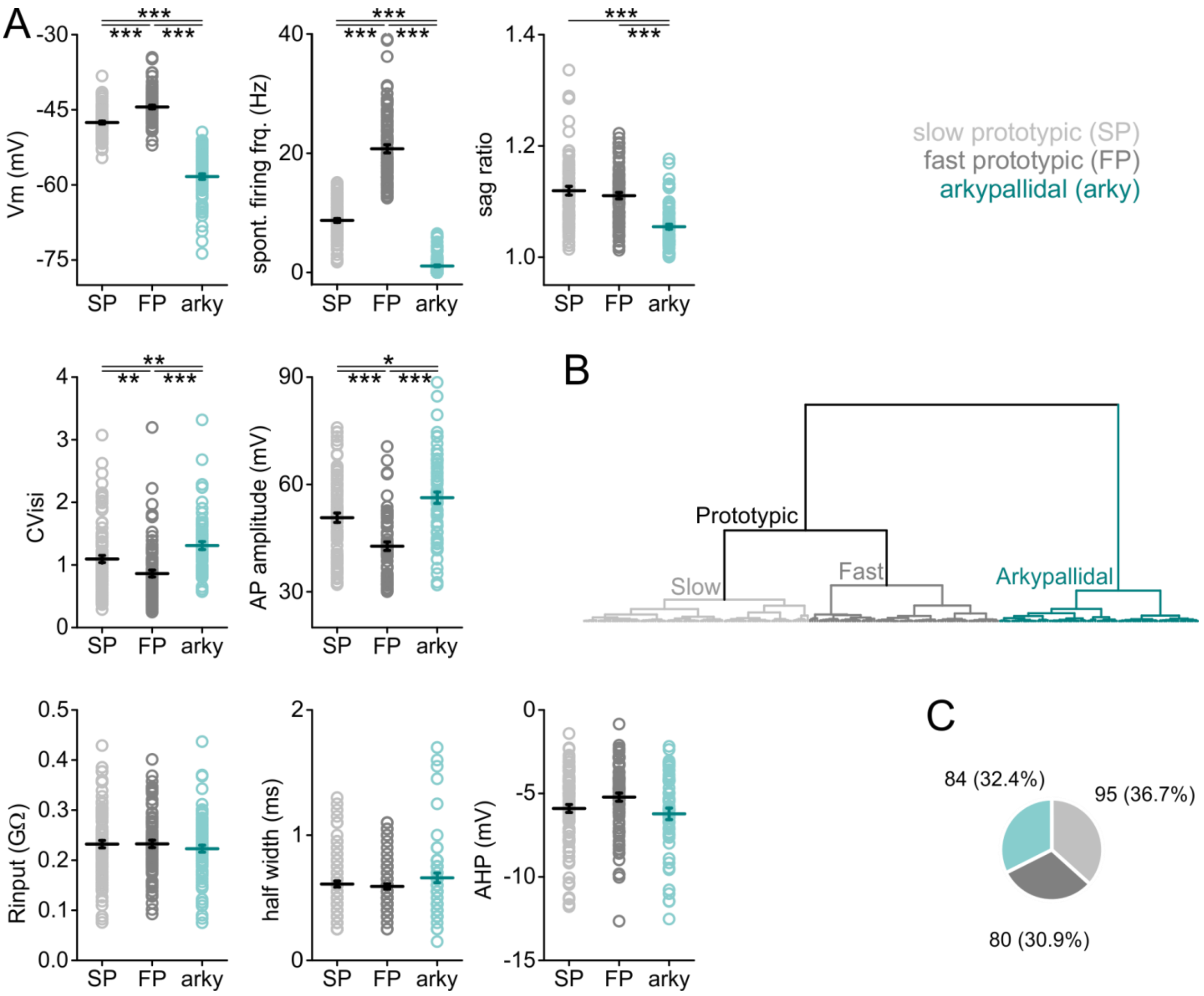


**Supplementary figure 2** (related to figure 2)**:** intrinsic properties of classified cells of the GPe divided into 3 groups. A. Intrinsic properties of the GPe cells according to their classification by membrane potential, spontaneous firing frequency and sag ratio. B. A dendrogram showing the classification of 259 recorded GPe cells. C. Pie chart of the cell numbers and proportions (bottom right). Slow prototypic cells in light grey, fast prototypic cells in dark grey, arkypallidal cells in turquoise, *** $p<0.05$, ** $p<0.01$, *** $p<0.001$. Data are presented as mean ± SEM. Statistical tests: Kruskal-Wallis ANOVA followed by Mann-Whitney test / one-way ANOVA with Tukey test mean comparisons.

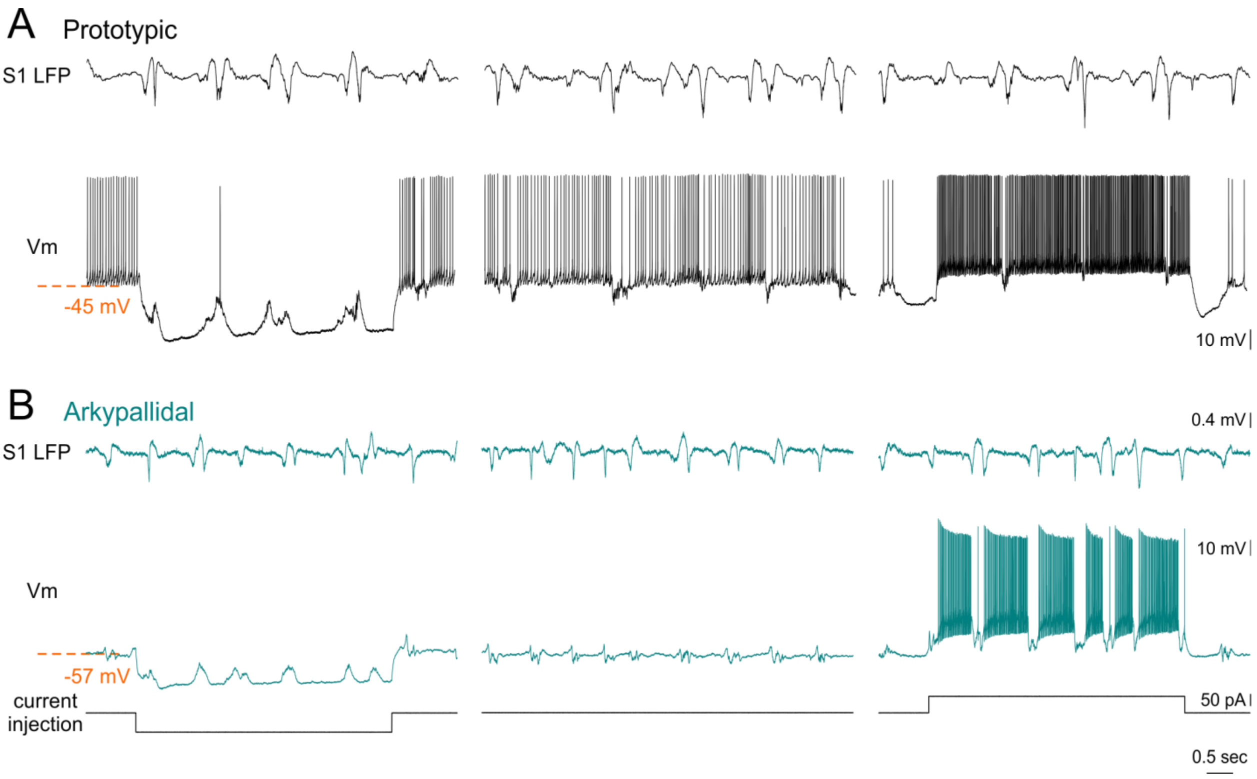


**Supplementary figure 3** (related to figures 1 and 3)**:** *In vivo* whole-cell recordings from prototypic and arkypallidal GPe cells during current injections. A and B *middle panel*. Prototypic cells (black, A) fire at high frequencies and slow or pause their firing when the cortex is engaged in an ‘up’ state, while arkypallidal cells (turquoise, B) are less active and often completely quiet. *Left panel*: Hyperpolarization exposes the ‘sag’ in both populations. During hyperpolarization, prototypic cells seem more similar to arkypallidal cells, depolarizing during up-states. *Right panel*: Depolarization increases the firing frequency of both populations and makes the arkypallidal cells more similar to prototypic cells, hyperpolarizing during up-states.

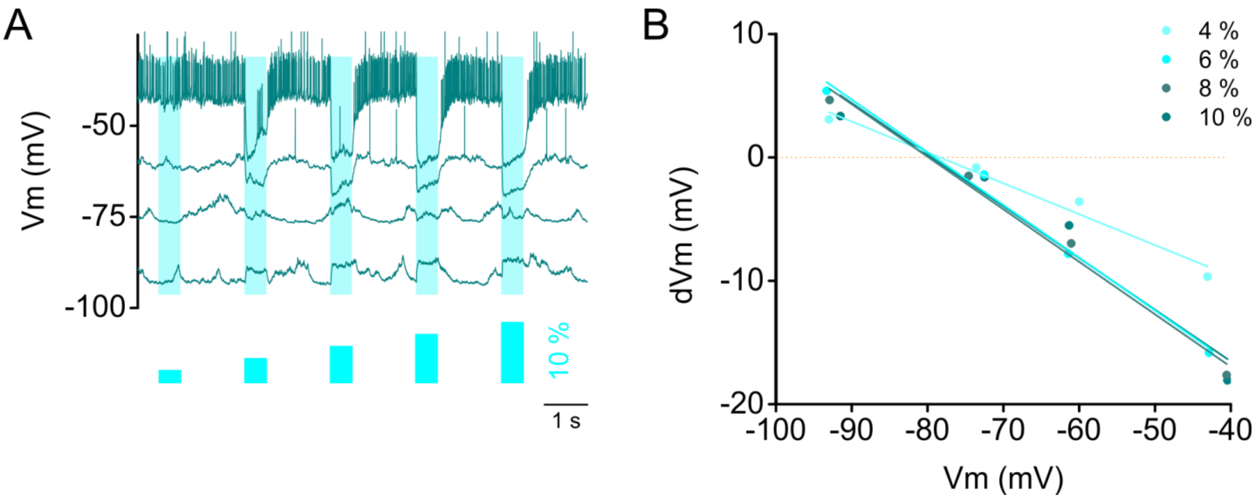


**Supplementary figure 4** (related to figure 4)**:** properties of the prototypic to arkypallidal inhibitory connection. To study the properties of the prototypic to arkypallidal inhibition we recorded the response in an arkypallidal cell to photostimulation of neighboring prototypic cells. A. responses in arkypallidal cell held at different membrane potentials using current injections. Areas shaded with light blue indicate the photostimulation times and relative intensities. B. response amplitude (dVm) as a function of holding membrane potential in 4 different light intensities (4 - 10% of maximum light intensity). The reversal potential of the inhibitory response corresponds to the chloride equilibrium potential of the pipette solution.

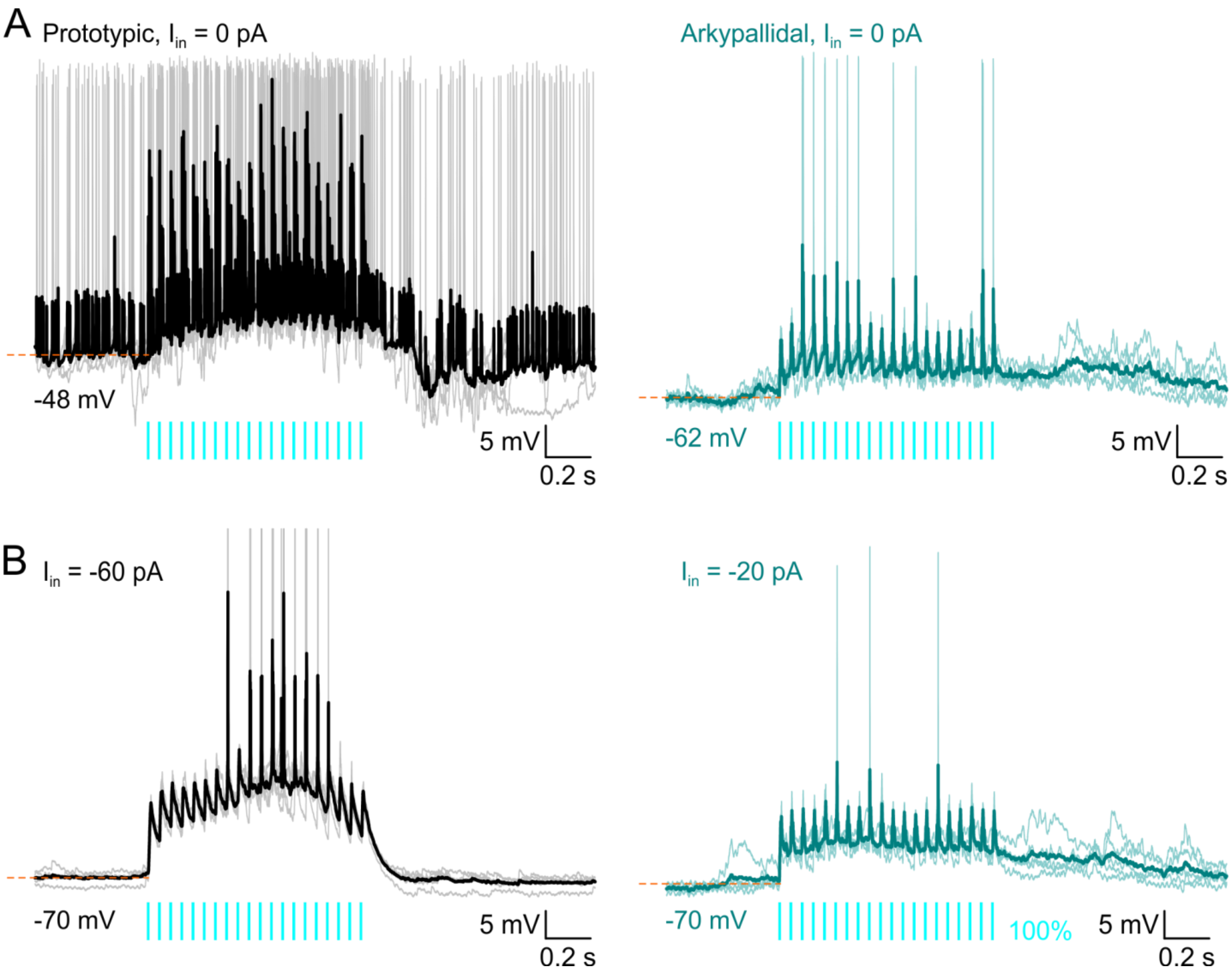


**Supplementary figure 5** (related to figure 5): 20 Hz STN photostimulation depolarizes and induces firing in both prototypic and arkypallidal cells. A. responses of prototypic (left) and arkypallidal (right) cells at rest to 20 Hz train photostimulation of the STN. Individual traces are presented in faint color superimposed with a thicker average trace. B. responses of prototypic (left) and arkypallidal (right) cells held at -70 mV with current injection to 20 Hz light stimulation of the STN. Individual traces are presented in faint color superimposed with a thicker average trace. Prototypic in black, arkypallidal in turquoise. Individual action potentials are clipped to maintain figure scaling.

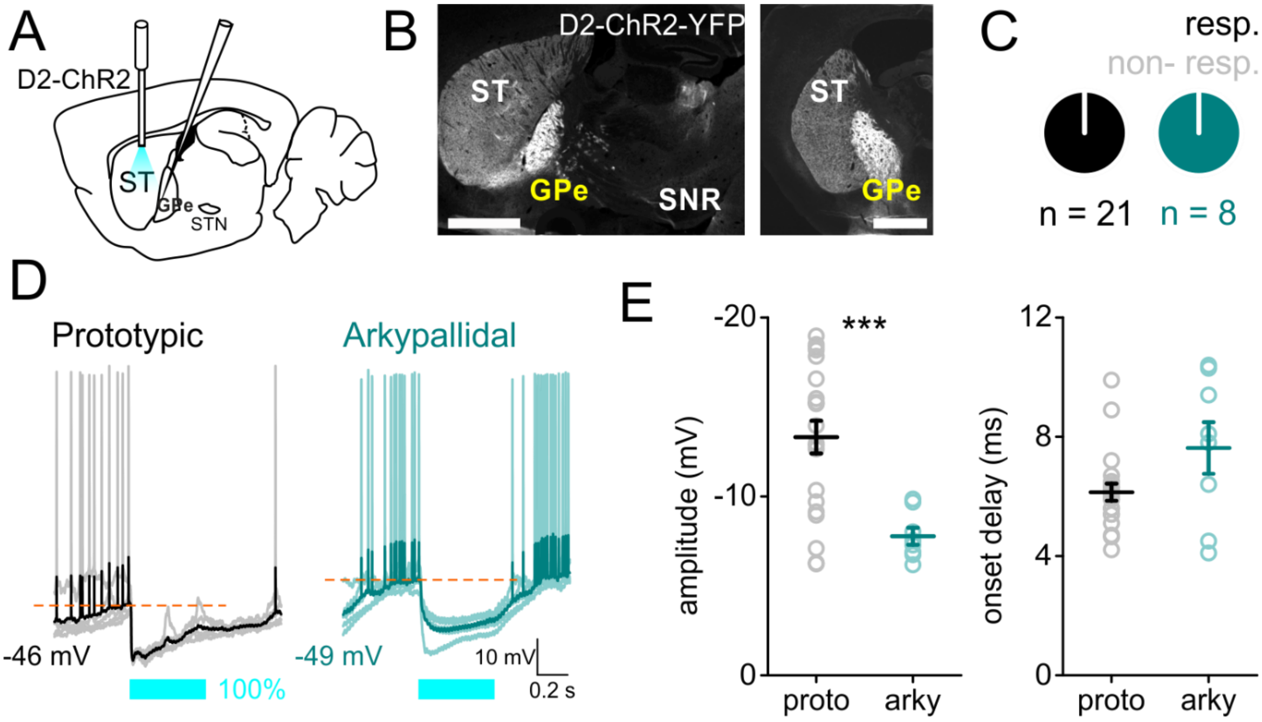


**Supplementary figure 6** (related to figure 6)**:** indirect-pathway MSNs target both prototypic and arkypallidal cells in D2-ChR2 transgenic animals. A. a scheme of the experimental set up, photostimulation and recording. B. sagittal (left) and coronal (right) sections showing the projection of indirect pathway MSNs to GPe. Scale bar 1 mm. C. pie chart representation of the proportion of the GPe cells responding to iMSN photostimulation. D. responses of prototypic (black, left) and arkypallidal cells (turquoise, right) to 500 ms photostimulation of striatal iMSNs. E. light response amplitude (left) and onset delay (right) in prototypic and arkypallidal cells. Prototypic in black, arkypallidal in turquoise. ST striatum, GPe external globus pallidus, SNR substantia nigra *pars reticulata*. Data are presented as mean ± SEM. Statistical tests: Mann-Whitney
